# Multichannel optogenetics combined with laminar recordings for ultra-controlled neuronal interrogation

**DOI:** 10.1101/2021.01.13.426504

**Authors:** David Eriksson, Artur Schneider, Anupriya Thirumalai, Mansour Alyahyay, Brice de la Crompe, Kirti Sharma, Patrick Ruther, Ilka Diester

**Affiliations:** Optophysiology, University of Freiburg, Faculty of Biology, 79104 Freiburg, Germany; Department of Microsystems Engineering (IMTEK), Cluster of Excellence BrainLinks-BrainTools, University of Freiburg, 79104 Freiburg, Germany; Optophysiology, University of Freiburg, Faculty of Biology, BrainLinks-BrainTools, Bernstein Center Freiburg, 79104 Freiburg, Germany

## Abstract

Simultaneous large-scale recordings and optogenetic interventions hold the promise to decipher the fast-paced and multifaceted dialogue between neurons that sustains brain function. Here we developed unprecedentedly thin, cell-sized Lambertian side-emitting optical fibers and combined them with silicon probes to achieve high quality recordings and ultrafast multichannel optogenetic inhibition in freely moving animals. Our new framework paves the way for large-scale photo tagging and controlled interrogation of rapid neuronal communication in any combination of brain areas.

**Highlights:** To combine large-scale recordings with optical perturbation we have developed a number of new techniques such as thin optical side-emitting fibers, a fiber matrix connector for thin fibers, an electro-optical commutator for multiple thin fibers, an active patch cord, a flexible fiber bundle ribbon cable, and a modular multi-optrode implantation holder.

## Main

Combined optogenetic manipulations and elecrophysiological recordings have been pioneered with a two-tipped optrode ^1^ and optotetrodes, i.e. an optical fiber surrounded by tetrodes ^2^. This has been refined with a single-tip approach ^3^, integrated fibers in electrode arrays ^4^ as well as using micro light-emitting diodes (μ-LEDs) combined with recording probes or co-integrated in silicon-based probes ^5–8^. Here we focused on the optimization of optical fibers as they allow targeting single cells as well as well as covering large volumes while keeping the flexibility to apply any desired wavelength via an external exchangeable light source. The existing fiber-based approaches illuminate the tissue surrounding the electrode either from a fiber at the top of the electrode shank^9–11^ or from a tapered fiber at some distance in front of the electrode shank ^12^. Although the elegant tapered fibers produce a more even illumination along the electrode shank, the light is typically strongest towards the fiber tip ^13,14^. Furthermore, the relatively stiff large-diameter fibers render the fiber location in the tissue dependent on the location of the connector ^15^, and for multiple fibers it may become difficult to avoid rupturing the vasculature. Moreover, multifiber stimulation would require a multichannel optical commutator for long-term stimulations in the freely moving animal. Finally, it is a common problem that optical stimulation next to extracellular electrodes causes photoelectrochemical, electromagnetic interference and photovoltaic effects ^16,17^, not to mention the mechanical damage induced especially by the larger fibers^18,19^ which hamper the recording quality.

Here we propose an alternative optrode design for silicon probes based on multiple ultrathin optical fibers which we call Fused Fiber Light Emission and eXtracellular Recordings (FFLEXR). We mitigate the above-mentioned issues with thin linearly emitting optical fibers that can be attached to any silicon probe, a lightweight fiber matrix connector, a flexible fiber bundle ribbon cable, an optical commutator for efficient multichannel stimulation, active patch-chord, and a Photo Voltaic Response (PVR) management algorithm.

To allow simultaneous extracellular recordings and multichannel fiber targeting in the freely moving animal we developed an optical commutator that was integrated into an electrical commutator (**Fig. 1A**). Light from a laser was guided to one of four patch cord fibers via a galvo scanner and a scan lens. The locations of the fibers were determined by scanning the optical ferrule which channels the light into the patch cord, and measuring the transmitted light with a linear intensity meter clip that was attached to the animal patch cord (**Fig. 1A** and **SFig. 1A-B**). Optical fibers with a distance of 100 μm can be addressed individually with a crosstalk of 0.3%, leading to a maximum of 100 individually addressable fibers per square millimeter in the ferrule (**SFig. 1C**). We were able to couple 49.3±7.3% of the input power into a 60 μm fiber with a 50 μm core and 14.3±6.6% for a 30 μm fiber with a core of 24 μm (**SFig. 1D**). An angular encoder allowed the galvo scanner to track one of the selected fibers as the animal, the motorized commutator and the ferrule rotated. The average mechanical error of the optical commutator was 3.3 μm (**SFig. 1E**).

**Figure 1.**
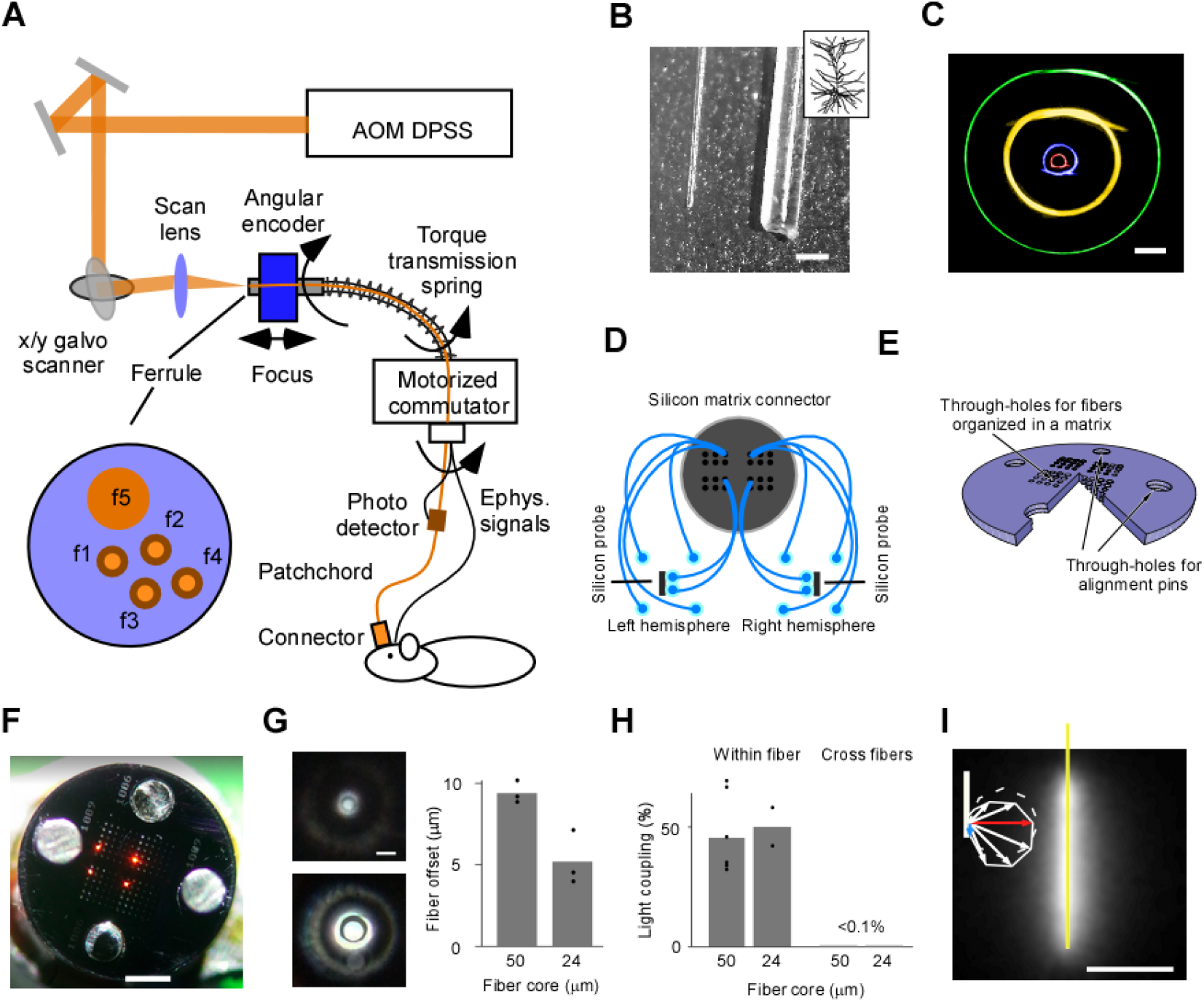
Optical framework for applications of ultrathin fibers with shaped pattern write-in in the freely moving animal. A: Optical system for automatic fiber bundle detection and optimized fiber coupling from an Accusto Optically Modulated (AOM) Diode Pumped Solid State (DPSS) laser. Feedforward control from the ferrule angle allows the galvo scanner to track one of the selected fibers in a freely moving animal. B: A 30 μm fiber next to a 230 μm fiber. Scale bar, 200 μm. Inset depicts a schematic of a neuron to illustrate the size relationship. C: Minimal bending radius for fibers with different outer/core diameter: 230/200 μm (yellow), 125/100 μm (green), 65/50 μm (blue), and 30/24 μm (red). This flexibility is crucial for flexible ribbon cables and for dissociating the location of the fiber end from the position of the fiber matrix connector. Scale bar, 2 mm. D: Spatial relation between implantation location and fiber matrix connector. E: Simplified 3D schematic of the circular fiber matrix connector plates with a triangular cut-out to illustrate the cross section through the 380-μm-thick plate realized in silicon using microsystems processing (four 4×4 array illustration for improved visibility, more fibers are possible). F: Photograph of fiber matrix connector. Fibers were reversely illuminated to visualize their location in the fiber matrix connector. Scale bar, 1 mm. G: Left: Fiber hole alignment for a 30/24 μm outer/core diameter fiber (top) and a 65/50 μm outer/core diameter fiber (bottom). Scale bar, 50 μm. Right: Quantification of the offset between two connector plates for two different holes sizes for 30 μm diameter fiber (24 μm core) and for 65 μm diameter fiber (50 μm core). H: Connector transmission efficiency and crosstalk between neighboring fibers in the connector. I: A coated 2 mm fiber in milk. Scale bar, 1 mm. Note that the radial side-emitted component (red arrow) is larger than the axial forward component (blue arrow).

To achieve high quality extracellular recordings next to large-scale optical interventions, we implanted optical fibers with an outer and core diameter of 30 μm and 24 μm, respectively (**Fig. 1B**). In comparison to fibers with a larger outer/core diameter, the thin 30 μm fiber was very flexible (**Fig. 1C**), thus allowing the implantation location to be independent of the fiber connector location (**Fig. 1D**). The fibers in the patch cord had a core diameter of 50 μm and each such fiber was targeting two or four 30 μm-fibers in the animal.

The fiber matrix connector was manufactured in silicon using photolithography and deep reactive ion etching (DRIE) (**Fig. 1E** and **SFig 2A-B**). Using 1 mm diameter precision guide pins (**SFig. 2C**), with 0.995 mm diameter lower boundary, the alignment of two fibers connection plates was 5.2±1.7 μm for the 30 μm fiber and 9.4±0.7 μm for the 65 μm fiber, and the light transmission efficiency was in the order of 50±11% for the 30 μm fiber and 45±15% for the 65 μm fiber, with a minimal cross talk between neighboring fibers (**Fig. 1H**).

We manufactured linearly emitting fibers from 30 μm fibers by manually polishing them with a rotating 5 μm diamond paper (**SFig. 3A**). The result was a linearly emitting fiber with a stronger axial (rotational symmetric and forward directed) component than radial (rotational symmetric and sideward directed) component (**SFig. 3Bi, ii** and **C**). To approach Lambertian emission, with a stronger radial than forward component, we coated the fiber with a thin diffusing layer (**Fig. 1I** and **SFig. 3Biii, iv** and **D**). A large range of emission lengths (0.5-10 mm) can be manufactured using this simple procedure (**SFig. 3E-G**).

To manufacture the implant, two uncoated- or two coated side-emitting fibers were attached immediately at the back-side (i.e. facing away from the electrode contacts) of a 75 μm wide silicon probe. Four additional fibers (from now on called fiber matrix) surrounded this back emission (BE) probe (**SFig. 4A**). The same configuration was used for the other hemisphere. BE-probes, fiber matrix, light connector and two electrode connectors were temporarily supported by a 3D printer holder during implantation (**SFig. 4B**). The total weight of the finalized connector with an epoxy filled 3D printed connector sleeve (**SFig. 4B**) was 0.25 g. Implantation was guided by the in vivo fluorescence (**SFig. 4C**). The 4 mm long fibers did not bend during implantation and the dimpling of the brain was minimal when those BE-probes and fibers were implanted (Supplementary video).

To demonstrate the viability of our approach in freely moving animals, we implanted 7 rats with combinations of the BE-probe and the fiber matrix. We recorded electrophysiological signals and applied yellow laser light (**SFig. 5A**) in freely moving rats in sessions of 60 to 120 minutes over the course of 10 days. To test the light distribution along the optical fiber, we took advantage of the PVR amplitude (**SFig. 5B**) along the electrode shank (**SFig. 5C**). For the uncoated side-emitting fiber there was a significantly larger PVR towards the tip of the electrode (p=0.035, n=4, student T-test, slope: −0.96±0.52mm^-1^), while for the coated fiber the difference was not significant (p=0.57, n=5, student T-test, slope: −0.27±0.96mm^-1^, **SFig. 5D** and **E**). This illustrates that despite an even emission along the fiber, a Lambertian angular distribution is crucial for an even illumination intensity in a partially scattering and absorbing tissue. The strong scattering in the brain allows the light emitted from our side-emitting fibers, from the backside of the electrode, to go around the electrode shank such that the light reaches the electrodes with an intensity that is 39.8% of that at the electrodes (**SFig. 5F**). Moreover, the PVR generated by the BE probe corresponded to a light intensity that was 2.3 times that of the fiber matrix (**SFig. 5G**).

After 10 days of implantation, there were clearly discernible action potentials for both constellations with coated and uncoated fibers (**SFig. 5H**). The action potential amplitudes ranged between 100 and 200 μV with on average 0.77 sorted units per electrode (uncoated: 0.78±0.27, mean±sd, n=12 sessions, coated: 0.76±0.41, mean±sd, n=7 sessions) (**SFig. 5I**). In accordance with the equal recording quality, the astroglia staining decreased as a function of the distance to the fiber similarly for the coated and uncoated fiber (**SFig. 5K** and **K**). The BE-probe can thus support homogeneous illumination, and high-quality extracellular recordings.

The final prerequisite for robustly aligned recordings and stimulations is the handling of light artifacts, such that extracellular action potentials can be reliably detected before, during, and after optogenetic inhibition. In 3 rats that had been injected with AAV2/5 carrying a hSyn-eNpHR3.0-mCherry construct we recorded neural activity with the BE-probe (**Fig. 2A**). Using a PVR management procedure (see methods), that can be applied after any spike sorting scheme, we demonstrate a drastic reduction of the influence of PVR on spike count statistics (**Fig. 2B**).

**Figure 2.**
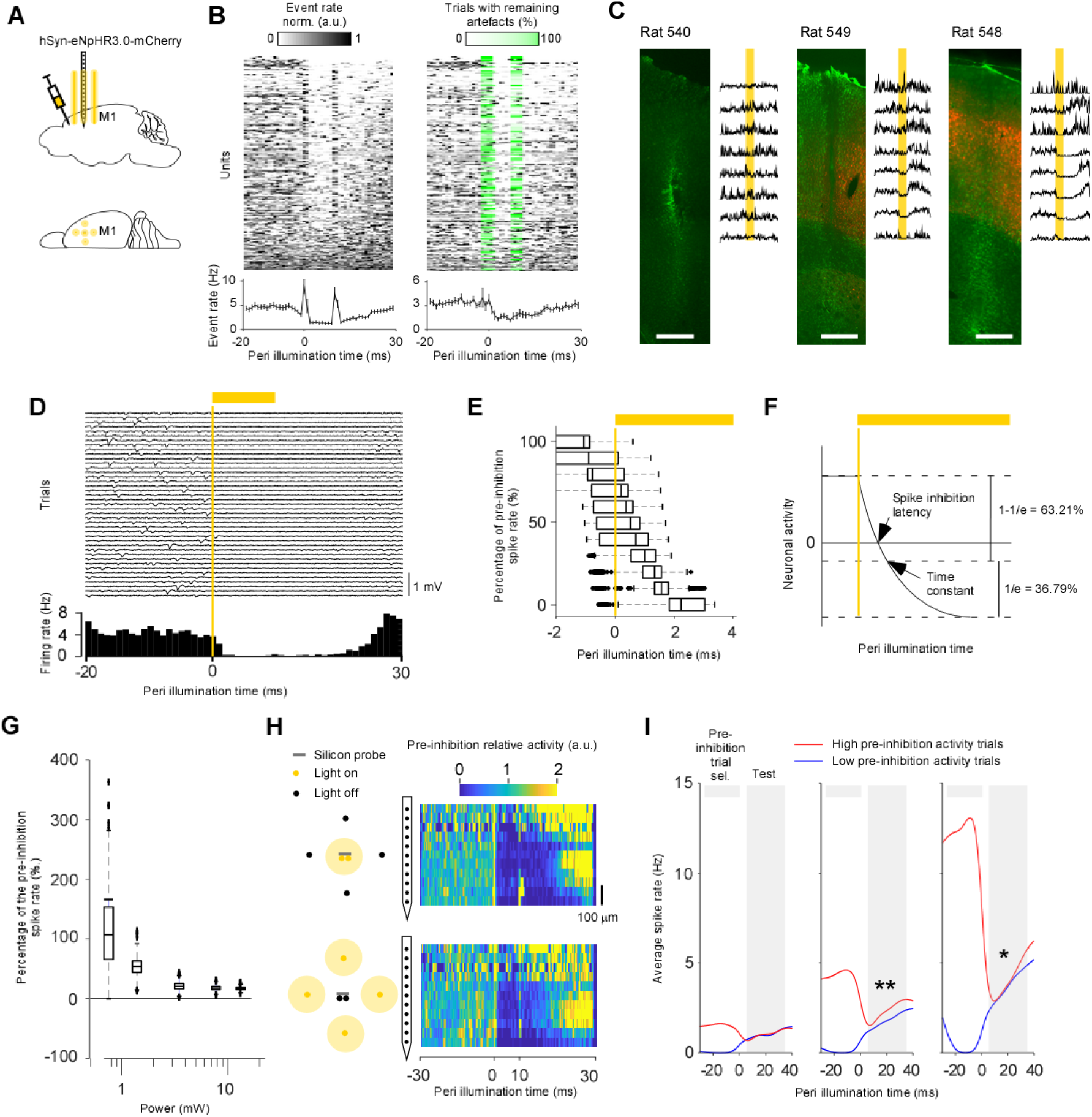
Ultrafast optogenetic inhibition in the freely moving animal. A: Injection, stimulation and recordings in M1. B: Activity of sorted units without (left) and with (middle) PVR management. The PVR management algorithm first subtracts an estimation of the PVR and then estimates in which trials the subtraction was incomplete. This remaining PVR trial percentage (green) is then used to weight the average across trials for each time point (lower left and right panel). Data is from animal 540, 548, and 549. C: Relation between opsin expression (red) and inhibition strength. To quantify the overall extracellular response, spikes were detected at a threshold of −30 μV on a, per channel basis without PVR management (PVR removed for visibility). GFAP immunostaining was used to identify the BE-probe location (green). Scale bar, 500 μm. D: Single trial extracellular traces for 40 inhibitions from one electrode channel (upper panel). Histogram of the spikes detected with a threshold of −30 μV (lower panel). E: Average firing rate across all units for animals 548 and 549. Error bars denote the standard error across units for each 1 ms bin. Latency at different percentages of the pre-inhibition spike rate bootstrapped across all sorted units. F: Illustration showing that the spike inhibition latency can be shorter than the time constant of the hyperpolarizing function. G: Percentage of the pre-inhibition spike rate as a function of the light power. The light power is the estimated total light power that exits the entire 2 mm side-emitting fiber. It is estimated from the intensity given by the active patch cord multiplied by the efficiency of the fiber matrix connector. H: Comparing optogenetic inhibition effects on neuronal responses using two constellations: BE-probe (top) and fiber matrix (bottom). To quantify the overall extracellular activity, spikes were detected at a threshold of −30 μV on a per channel basis without PVR management. I: Inferring neuronal activity after inhibition onset from pre-inhibition activity. The units were divided into three groups: low (left), moderate (middle), and high (right) average firing rate.

Next, we analyzed the neuronal responses to laser light. We found clear optogenetic inhibition in probes that passed through tissue that contained eNpHR3.0-mCherry expressing neurons (**Fig. 2C**). The buildup of inhibition was very quick (**Fig. 2D**) and completed within 2.2 ms (**Fig. 2E**). This is faster than the time constant of halorhodpsin ^20^. A strong hyper polarization will cancel spikes before the time constant has been reached (**Fig. 2E**), and several times faster than the more physiological, channel based or indirect inhibition ^10^, and reinforces the fundamental property of optogenetics in terms of a fast silencing of genetically and anatomically defined neurons ^21^. The low latency inhibition of genetically defined neurons is a prerequisite for reliable photo tagging (**Fig. 2D** and **E**). Spikes were typically eliminated for the duration of the light pulse, or longer. The prolonged suppression of spikes outlasting the actual inhibition time has been described for optogenetic inhibition ^22,23^ and for physiological inhibition ^24^. The inhibition strength reached an asymptote at around 3 mW, corresponding to a power density of 12.5mW/mm^2^(**Fig. 2G**). The inhibition was similar for BE-probe and fiber matrix emission (**Fig. 2H**), which confirms similar PVR amplitudes for both conditions (**SFig. 5F**). Finally, it is conceivable that the pre-perturbation activity of the perturbed neurons can affect the activity during or after the inhibition ^25^. To this end we divided the trials in high and low preinhibition spike count for three different groups of neurons: low, moderate, and high firing rate neurons (**Fig. 2I**). In general, trials with a high pre-inhibition activity had a higher activity after inhibition onset and this effect was significant for neurons with moderate and high firing rates (low rate: p=0.98, n=82, 0.0018±0.11 Hz, mean ± sem, Moderate rate: p=0.0035, n=82, 0.53±0.18 Hz, and high rate: p=0.025, n=82, 0.46±0.2 Hz).

The stimulation and recording framework presented here allows for maximally controlled interrogation of neural circuits since dense neuronal recordings can be conducted before, during and after optogenetic perturbations. Here we have enabled this by rendering fiber-based experiments compatible to, and as flexible as, extracellular recordings. This allows any desired combination of fibers and electrodes (**Fig. 3**). Since any spatial extent of the light emission of one fiber can range from 24 μm to 10 mm, and since a matrix of fibers can be used, it is possible to delineate brain areas of complex shapes. The flexible fiber bundle and the connector practically allow any combination of areas to be addressed, as well as individualized designs for each animal to avoid blood vessels and to target areas defined by intrinsic imaging, in vivo fluorescence, or computer tomography. The thinness of the fibers allow simultaneous extracellular recordings in all targeted areas to confirm the effectivity of optogenetic intervention and to control for changes in excitability ^26,27^, as well as to optically tag neuronal subtypes and to understand how trial by trial variability in the neural activity changes the context of perturbations of specific neuronal subtypes in freely moving animals.

**Figure 3.**
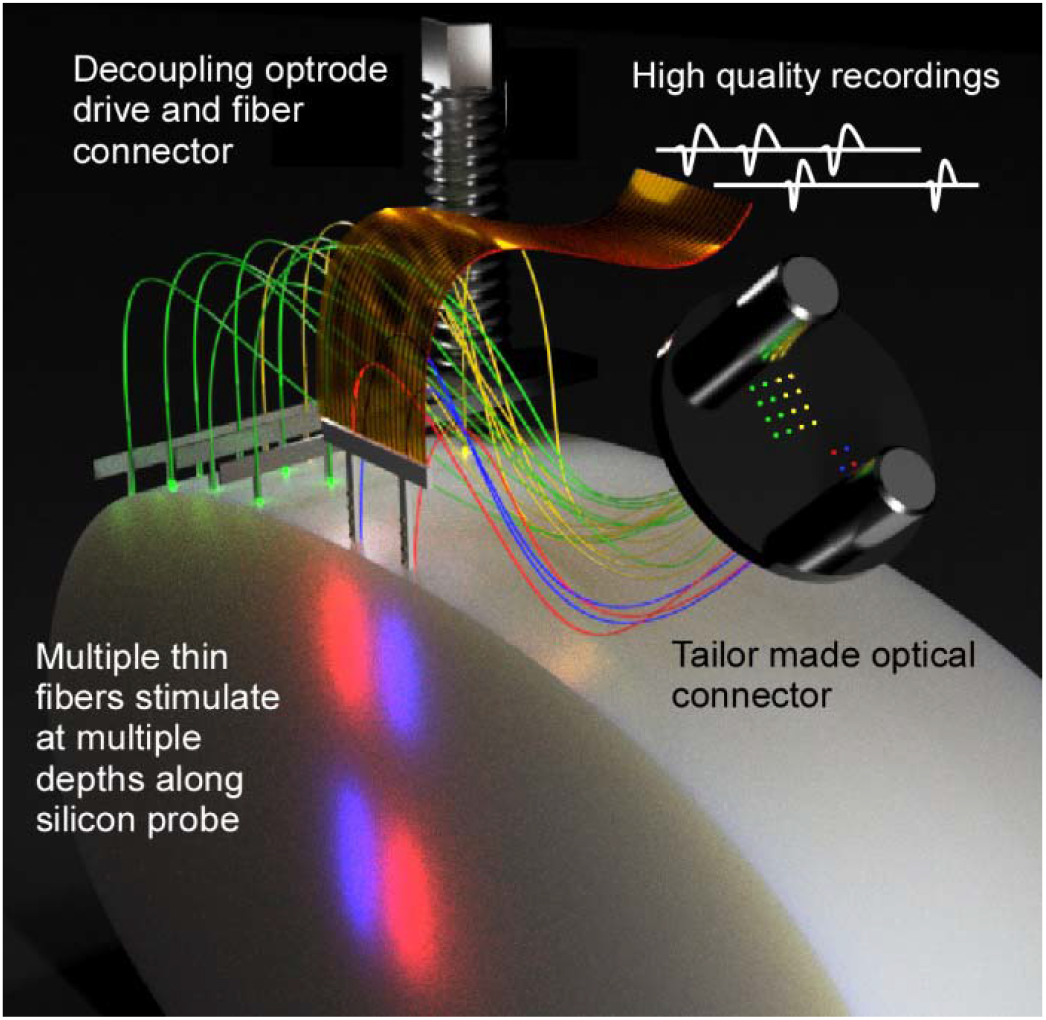
Schematic illustration of a possible fiber and electrode arrangements. The thin fibers allow both spatial flexibility and high-quality extracellular recordings.

## Methods

### Producing the fiber connector using MEMS technologies

The process performed in the cleanroom facility of the Department of Microsystems Engineering (IMTEK) of the University of Freiburg applied standard 4-inch, double side polished silicon (Si) wafers with a thickness of 380 μm. The wafer front side (FS) was coated in a first step with a 3.5-μm-thick silicon oxide (SiO_x_) layer realized using plasma enhanced chemical vapor deposition (PECVD). Next, UV photolithography applied a 10-μm-thick positive resist AZ9260 (Microchemicals GmbH, Ulm, Germany). The resist served as a masking layer in the subsequent reactive ion etch (RIE) process used to patterned the SiO_x_ layer on the wafer FS. Then, wafers were FS etched to a depth of 200 μm using the advanced Si etch process also known as Bosch process or deep reactive ion etching (DRIE) in an inductively couple plasma (ICP) etch system.

Next, the FS masking layer was removed by wet chemical etching using 5% hydrofluoric acid (HF). Wafers were then thermally oxidized to a layer thickness of 500 nm. This silicon dioxide (SiO_2_) layer conformally covered the wafer surface, i.e. the etched wafer front as well as the planar rear side. It served as an etch stop inside the recesses etched from the wafer FS. Then, a 2-μm-thick PECVD SiO_x_ layer was deposited on the wafer rear side (RS) serving as masking layer. It was patterned by UV lithography and RIE. Subsequently, DRIE was used to etch the wafers to a depth of ca. 150 μm. Then, the wafers were fixed on a handle wafer using a thermally conductive wax-like material, i.e. Cool Grease, used to keep the wafer temperature low and etched through with the DRIE process stopping at the thermally grown SiO_2_ layer. The handle wafer was removed by dissolving the Cool Grease layer. The circular connector plates were finally released from the wafer by application of torsional forces in order to break the suspension arms. The 500-nm-thin SiO_2_ layer inside the through-holes easily cracked during the subsequent cleaning steps and did not block the circular openings of the connector plate.

To accommodate for process tolerances, we implemented the large 1mm through-holes (for guide- and anchoring pins) with diameters between 994 μm and 1015 μm increased in steps of 3 μm (the large holes were made to fit guide pins with a maximal diameter of 1mm and a minimal diameter of 0.995). In the plate center, four identical square arrays with 6×6 smaller circular through-holes are arranged in the middle of the connector plate. Within an array, the through-holes are positioned at a center-to-center distance of 150 μm. The through-holes on the wafer front side (FS) had diameters varying from 22 to 39 μm, in steps of 1 μm, and 49 to 83 μm in steps of 1 and 2 μm, respectively. The diameter on the wafer rear side (RS) was enlarged by 10 μm each to compensate for a potential misalignment between the FS and RS lithography process steps described below. In case of the larger though-holes with a diameter of 1 mm used for the alignment pins, we added disk-shaped dummy structures. Those were used to maintain across the entire wafer similar lateral dimensions of the structures to be etched. This feature has been implemented for achieving comparable etch rates of the smaller fiber though-holes and the larger trenches defining the disk shape and the large through-holes.

### General fiber handling

Since the 30 μm fibers are practically invisible on a white background, the workshop desk was covered with a black poster board (TB5, Thorlabs, Germany, Bergkirchen). To handle the fibers we added a layer of Kwik Cast (WPI, Germany, Friedberg) to the grabbing surfaces at the tip of a forceps. Finally, since it is difficult to pick up a fiber lying on a flat surface with the bare fingers, we used a sticky stick to pick up fibers. This sticky stick was made by squeezing a small amount (~0.1ml) of Paddex (UHU, Germany, Bühl/Baden) through a 1 mL syringe such that a small amount of sticky clay protruded from the hole in the tip of the syringe.

### Polishing side-emitting fibers

The side-emitting fibers were created by gradually polishing away the cladding from a thin optical fiber (30μm cladding, 24μm core, NA=0.86, S17, Lifatec, Germany). To evaluate the polishing, the other end of the fiber was attached to a light source. The cladding was removed using a 6 μm diamond polishing paper (LF6D, Thorlabs, Germany, Bergkirchen). The polishing paper was attached to a disc rotating with 10 turns per second with a diameter of 60 mm. The fiber was gently pushed towards the rotating polishing paper using a cotton tip for which the wooden end was covered by a rubber coating (Kwik Cast, WPI, Germany, Friedberg). To polish different sides of the fiber it was fixed inside a tube which in turn could be rotated. After the polishing the light was escaping the fiber with a higher intensity to the front than to the back. A true side-emitting fiber with equal intensity to the front and back and maximal intensity to the sides was achieved by applying a thin diffusing layer of nail polish diluted by 1:4 with acetone (NailPolish Misslyn, nail polish, white, 90, Innovative Cosmetic Brands, Karlsfeld, Germany). Once the fiber was finished it was cut to a length suitable for handling (~80 mm). In a serial fashion, additional fibers could be made by pulling out the fiber from the rotating tube since the total fiber length was around 1 m.

### Assembling the implant

The implant was assembled on a 3D printed substrate. This substrate served to hold fibers, electrodes and their connectors during implantation with a single stereotactic holder. As the implantation proceeded parts of the substrate was melted away with a standard cauterizer (Bovie, WPI, Germany, Friedberg). The 3D printed substrate had a two-step staircase. On the longest staircase, the electrodes and fibers were located that should be implanted first. Once those had been implanted their plastic could be removed and the electrodes and the fibers at the second staircase could be implanted.

Fibers, electrodes, and their connectors were attached to the substrate as follows. To hold the fibers and BE-probes for one hemisphere (one staircase), a honeycombed arrangement of 19 guide tubes (i.e. two guide tube layers away from the center guide tube), with an inner diameter of 200 μm and an outer diameter of 350 μm (TSP200350, Polymicro, Arizona, Phoenix), with a length of 3 mm, were glued to each staircase. The BE-probes consisted of two side-emitting fibers attached on the backside of the laminar electrodes (two 30 μm diameter fibers fit behind a 75 μm wide electrode). The fibers were attached to the electrodes using super-glue (UHU, Germany, Bühl/Baden) at the base of the laminar electrodes. Then the BE-probes were inserted in the middle guide tube on respective staircase and four fibers were inserted in four of the most peripheral guide tubes at a distance of 600 μm from the BE-probe in a star-like fashion. To protect the fibers during implantation, and to form a fiber bundle ribbon cable, they were coated with rubber up to a distance of 10 mm from the guide tubes.

Before, the fibers were inserted in the silicon plate we added two 1 mm diameter and 3 mm long low-precision steel pins in the guide pin holes that were not used for the high precision pins. This ensures a rigid connection between the silicon plate and the rest of the connector. The four fibers on respective staircase was inserted into one 65 μm hole. The two fibers coming from one BE-probe was put into another 65 μm hole and two additional dummy fibers were added. The dummy fibers prevent unnecessary heat buildup at the connector junction. The fiber length between silicon plate and guide tubes was set to 35 mm. The same procedure was repeated for the other staircase/guide tube bundle. The fibers were attached to the silicon plate using heat curable epoxy. To this end small drops of epoxy (using a 33 Gauge wire) was added to all the fibers from both sides of the plate to prevent undercutting during polishing. A small heat coil with a diameter of 1 mm and a length of 5 mm (200 μm diameter Nichrome wire with a current control to achieve an orange glowing color) attached to a pen like grip was used to heat up the air in the vicinity of the epoxy. A heat gun could not be used since that would blow away the small parts. Before the plate was polished the remaining bare 25 mm fibers was coated with rubber to finalize the fiber bundle ribbon cable. A standard surgical stereo microscope with 40 times magnification was used to inspect the polishing.

Next the guide pins should be added to the silicon plate. An indent was polished in one end of the guide pins to aid their fixation in the contact. The other end was inserted into a fixture that ensured the right spacing and a 90° angle of the guide pins. The fixture with the two guide pins was coated with Vaseline to prevent epoxy from clogging the fixture. This assembly was coupled to the polished silicon plate with the fibers. A 3D printed connector sleeve was slid over the rubber coated fibers and down to the contact with the guide pins. Finally, the connectors sleeve, fibers, silicon plate and guide pins were epoxy together using slow curing epoxy (End-fest, UHU, Germany, Bühl/Baden).

### Assembling the active patch cord

The active patch cord relies on a small amount of light that escapes the fiber and passes through the protective tube to be detected by a linear photodetector on the outside of the patch cord. To this end we used a yellow tube with the inner diameter of 600 μm and an outer diameter of 900 μm (FT900Y, Thorlabs, Germany, Bergkirchen).

The assembly of the patch cord began with inserting the four fibers (AS50/60IRPI, Coating 65mm polyimide, LEONI Fiber Optics, Germany, Schierschnitz) into the protective tube. Then the fibers were gently inserted into the four holes of the silicon plate that corresponded to the holes used by the implant. For stability, two 1 mm diameter and two? 3 mm long low precision steel pins were inserted in the guide pin holes that were not used for the high precision pins. Before securing with heat curing epoxy, we ensured that all fibers had the same length before entering the protective tube. The identical fiber lengths prevent individual fibers to form knots or loops as the protective tube was slid towards the silicon plate. Next the silicon plate was polished and coupled to two high precision guide pins that were inserted in a Vaseline covered fixture. Two 3.5 mm tubes were cut from a 17 gauge injection needle. Those were filled with Vaseline and slid over the high precision guide pins that were extended from the patch cord side. Then a 3D printed patch cord sleeve was glued to the tubes, silicon plate, anchor pins, and tubes. The tubes added robustness to the connector. We used the same patch cord for all sessions in the freely moving rats and the power transmission for the patch cord was measured after all recordings were done (End-fest, UHU, Germany, Bühl/Baden).

The fibers on the other side of the patch cord were coated with heat curing epoxy and inserted into four guide tubes with an inner diameter of 70 μm and an outer diameter of 200 μm (TSP075200, Polymicro, Molex, USA). This bundle was inserted in a metal ferrule with an outer diameter of 2 mm that was filled with heat curing epoxy. Note that the arrangement of individual fibers is flexible since each fiber is addressed by the galvo scanner. Finally, the metal tube was gently heated to facilitate an even distribution of epoxy in the metal tube. Care was taken to slowly heat up the epoxy (such that the curing would emerge after around 20 seconds) to prevent heat induced cavities formed between the fibers which in turn would hamper proper polishing.

### Assembling the laser system

The laser system (see **Fig. 1A**) was designed to achieve high-intensity multichannel pulsed optogenetic inhibition. A high intensity laser source is favorable as it can compensate for the relatively large coupling losses that occur with small optical fibers. Here we used one 100mW DPSS bragg modulated laser (594nm, Mambo, Cobolt, Sweden, Solna). The side-emitting fibers spread out the photon current over a larger area thus preventing optical burning of the tissue. This in turn opens up for a more efficient optogenetic inhibition. Further, the 10 ms light pulses used here causes a minimal heat buildup.

The optical axis passed through a galvo scanner with two mirrors (8315KSM40B, Cambridge Technology, USA, Massachusetts, Bedford). The galvo mirror had a high-power servo amplifier unit and deflected the beam into one of the fibers within 100 μs. From the galvo mirror the light passed through a scan lens (LSM03_VIS, Thorlabs, USA) which in turn focused the light into one of the fibers in the patch cord. The ferrule of the patch cord was located in an angular encoder with 32,768 steps (7700D05000R00S0175N, Gurley Precision Instruments, USA, New York, Troy) that was supported by two ball bearings. A stepper motor stage (Mini25, Luigs Neumann, Germany, Ratingen) that could be moved along the optical axis served to focus the light at the ferrule in the optical commutator. The patch cord was turned by the motorized electrical commutator (MEC) (ACO32, TDT, USA, Florida, Alachua) when the animal turned and the angular encoder ensured that the galvo scanner always positioned the beam on the appropriate fiber. A metal spring transferred the torque from the motorized commutator to the angular encoder. In this way the optical axis of the laser system could be at a 90° angle to the rotating axis of the MEC. The patch cord passed through the MEC and down to the animal. The MEC, commutated the extracellular signals from 48 electrodes and two light intensity signals from the active patch cord. All signals were integrated via the QPIDe IO card (Quanser, Canada, Ontario, Markham) and Matlab scripting (Mathworks, USA, Massachusetts, Natick).

### Animals

All animal procedures were approved by the Regierungspräsidium Freiburg, Germany. In this study we used seven male Sprague Dawley rats (400 g, Janvier) which were implanted at the age of eight weeks. Three to four animals were pair-housed in type 4 cages (1500U, IVC typ4, Tecniplast, Hohenpeißenberg, Germany) before implantation and the animals were single housed after the implantation in type 3 cages (1291H, IVC typ4, Tecniplast, Hohenpeißenberg, Germany) under a 12 h light dark cycle (dark period from 8 a.m. to 8 p.m., active period during which training and experiments happened).

### Animal surgery and virus injection

Animals were initially anesthetized with isoflurane inhalation followed by intra-peritoneal injection of 75 mg/kg Ketamine (Medistar, Holzwickede, Germany) and 50 mg/kg Medetomidin (Orion Pharma, Espoo, Finland). The animals were put into a transportation container covered with an opaque cloth to facilitate the anesthesia. Once the animals were anesthetized, they were positioned in a stereotaxic frame (David Kopf Instruments, Tujunga, CA, USA) and their body temperature was kept at 36-37 °C using a rectal thermometer and a heated blanket (FHC, Bowdoin, USA). The anesthesia of the animals was maintained with approximately 2% isoflurane and 0.5 l/min O2. For pre-surgery analgesia, we subcutaneously (s.c.) administered 0.05 mg/kg Buprenorphine (Selectavet Dr. Otto Fischer GmbH, Weyarn/Holzolling, Germany). Every other hour, the animals received a s.c. injection of 5 mL isotonic saline. Moisturizing ointment was applied to the eyes to prevent them from drying out (Bepanthen, Bayer HealthCare, Leverkusen, Germany). The skin was disinfected with Braunol (B. Braun Melsungen AG, Melsungen, Germany) and Kodan (Schülke, Norderstedt, Germany). To perform the craniotomy, the skin on the head was opened along a 2 cm long incision using a scalpel. The exposed bone was cleaned using a 3% peroxide solution. For virus injections craniotomies were drilled bilaterally extending from −1 to +3 mm in the anterior posterior direction and from +0.5 to +1 mm in the lateral medial direction relative to Bregma. We injected 1 μl viral vector (AAV-hSyn-eNpHR3.0-mCherry-WPRE, UNC Vector Core, North Carolina, Chapel Hill) at a rate of 100 nl/min into the respective subareas using a 10 μl gas-tight Hamilton syringe (World Precision Instruments, Sarasota, FL, USA). To minimize the reflux of the injected volume, we left the injection needle in the tissue for 10 additional minutes before slowly extracting it out of the brain. To protect the brain during virus expression the craniotomy was covered with bone wax. The surgical site was closed around the implant by interrupted sutures using 5-0 silk (SMI AG, St. Vith, Belgium).

### Animal surgery and BE-probe implantation

The initial steps of BE-probe implantation were identical to those of the virus injections (see above). Self-tapping skull screws (J.I. Morris Company, Southbridge, MA, USA) for reference for extracellular recordings were placed over cerebellum. For BE-probe implantation craniotomies were drilled bilaterally extending from −2 to +5 mm in the anterior posterior direction and from +1 to +3 mm in the lateral medial direction relative to Bregma. The BE-probes with the surrounding fiber matrix were implanted bilaterally beginning with the right hemisphere. To this end, the degradable stair-case implant holder (see Assembling the implant) was held with a stereotaxic manipulator arm and positioned such that the lowest stair-step was located over the right hemisphere. The electrode- and fiber ribbon bundle was facing anteriorly and the 3D printed substrate was facing posteriorly. The right BE-probe shank was inserted 2 mm into the cortex while ensuring that no blood vessel was punctured by the BE-probe or the fiber matrix. Kwik-Cast (WPI, Sarasota, FL, USA) was applied to the craniotomy to protect the brain from the dental cement (Paladur, Kulzer GmbH, Hanau, Germany), that, in turn, was applied to anchor the first stair-step to the bone. Five minutes later, when the dental cement had cured, a cauterizer was used to cut the degradable implant holder just above the first stair-step in order dissociate the implant holder from the implanted stair-step. Next the remaining left stair-step was positioned over the left hemisphere by moving in the stereotaxic manipulator arm. The BE-probe was inserted to a depth of 2 mm, Kwik-Cast was applied and the stair-step was anchored to the bone. After five minutes the dental cement had cured and the left stair-step could be dissociated from the degradable holder. At this stage only the fiber matrix connector and zif-clip connectors were held by the stereotaxic manipulator arm. Those connectors were dissociated from the holder using the cauterizer and attached to the skull with dental cement. Finally, the fiber and electrode ribbon cables were protected with dental cement.

### Calculating the intensity ratio between the BE-probe and the fiber matrix

To calculate the light intensity ratio at the electrodes between the BE-probe-fibers and the fiber matrix, we equalized the PVR for the fiber matrix and BE-probe fibers by changing the relative intensity (given by the active patch cord) for those two fiber configurations. For example, by fixating the intensity for the fiber matrix we could modify the intensity for the BE-probe fibers such that the PVR became equal for the two configurations. Using the two resulting intensities we could calculate the ratio for the BE-probe and the fiber matrix. To find matching intensities we first estimated the range in PVR amplitudes for which they were linearly related to the light intensity. The relation between light intensity and PVR was linear up to around thousand microvolts, and for high light intensities the PVR saturated at +6000 μV. For each electrode, we estimated the time point after the laser onset for which the maximal intensity in the fiber matrix caused a PVR that was closest to thousand microvolts. If the largest PVR for a given channel was less than thousand microvolts, the time point of the maximal and the corresponding maximal PVR amplitude was used. This time point was used to look up the intensity of the BE-probe that generated the most similar PVR amplitude for the given channel. The intensity ratio was calculated as the maximal intensity for the fiber matrix divided by the latter intensity of the BE-probe.

### PVR management procedure

Since the light source was aligned with the electrodes, the Becquerel induced light artifacts were relatively large. The strategy here was to eliminate as much of the PVR as possible, and if a PVR could not be removed it was marked as “black out period”/“not a number” (NaN) in order to facilitate further processing. In short, first PVRs were decreased by subtracting PVR templates based on the applied light intensity. Second, a heuristic was used to estimate time points of remaining PVRs, time points that were “blacked out” for spike detection (the intensity of the green intervals in **Fig. 2A** denotes the percentage of trials with remaining PVRs). Finally, spike templates that were sorted during non-stimulated intervals, were fitted to threshold crossings during stimulation and were accepted as a true spike if the error was less than the standard deviation and if they did not contain a blackout interval. Thereby, the algorithm identified PVR waveforms and did not classify them as neuronal units.

To remove PVRs, data snippets were cut out beginning at 30 ms before the light pulse onset and ending 30 ms after the light pulse offset. Light onset and offset were defined according to the half maximum latency of the light intensity in the active patch cord. The resulting data snippets of 70 ms were classified into eight groups according to the four different fiber channels and the two different electrode shanks. The PVR removal was done individually for each of those eight groups. First the data was high-pass filtered. The data was super sampled by a factor of four to increase in the temporal resolution. Then the transient PVRs associated with light onset and offset were minimized. We tested several methods such as linear and nonlinear regression, principal component analysis, nonlinear principal component analysis and template matching. The best result was achieved with template matching. Due to the large number of pulses (in the order of 500 pulses) it was possible to tailor each template for each pulse. A template was created by averaging the 20 pulses with the smallest mean squared error to the pulse of interest. Averaging 20 pulses would result in a decrease of the signal amplitude (spike amplitude) of around 5%. Since the duration of each pulse varied, the templates for the onset and offset transient were calculated individually. To avoid discontinuities in the data, the templates for the onset and offset were calculated with an overlap of 3.3 ms (100 samples at 30000 samples per second) between the onset and offset. A linear interpolation was used to join the onset and offset template across this 3.3 ms (100 samples) overlap.

To detect remaining PVRs we applied a multistep procedure. To capture the high frequency remaining of the PVR, we derived the signal in time. The PVR remaining was concentrated where the PVR had its strongest temporal derivative, which was at around 260 μs after laser onset and lasted for another 300 μs. Since the error from the PVR removal can be both positive and negative we rectified the derived signal. To improve the PVR estimate we averaged the rectified signal across trials as follows. Since, the success of the PVR removal depended on the laser intensity, the trials were sorted according to the light intensity given by the active patch cord, and a moving window filter was run across the trials for each time point. The moving window filter averaged the rectified values across seven trials with the exception of the trial that had the largest rectified value. The largest rectified value was left out to minimize the influence of spikes. If the resulting signal for a certain trial and for a certain time point was larger than four times the standard deviation of the prestimulation period, it was regarded as a PVR and the corresponding data point in the PVR minimized data was replaced by NaN. This data was referred to as the result of a PVR management procedure.

Since it is unclear how KiloSort ^16^(or any other spike sorting routine) handles optical PVRs, we choose to run the spike sorting on the PVR free periods between the pulses by concatenating those periods to one matrix. The number of clusters was assumed to be two times the number of channels. After the spike sorting we run the automatic curation. Sorted units having a negative spike amplitude larger than 400 μV and three spike amplitude larger than 20 μV were omitted due to their unphysiological properties. The remaining sorted units were matched to spike events in the PVR managed data. This was done as follows. For a given trial all the channels were normalized to have standard deviations equal to one. The channels were sorted according of the linear layout of the probe. Then the smallest negative event was chosen (and marked with a NaN to not be chosen again) and compared to the waveforms of the sorted units. To this end, three channels centered on the negative event were subtracted from the corresponding channels of one of the waveform of the sorted units. The unit giving the smallest squared error was selected. If this error was smaller than one (the standard deviation of the data) the waveform was classified as action potential. After this, the procedure was repeated for the next spike. The procedure stopped when no event was left which was larger than four times the standard deviation.

To minimize a PVR related spike bias, an event detected in the spike matching procedure was regarded as invalid if the corresponding spike window (−20 to 40 samples surrounding the event) contained a NaN. A separate data structure was used to keep track of NaN’s. After the spike matching procedure the regions containing NaN’s were expanded 40 samples back in time and 20 samples forward in time. This served to label all periods in which no action potential could occur (because the spike window is not allowed to contain a NaN).

### Histology

Brains transferred from sucrose solution were attached to the cooling block of a microtome with Tissue Tek (Sakura Finetek, Fisher Scientific, Germany). The slices were transferred to vials containing Phosphate-Buffered Saline (PBS) with 0.01% sodium azide. For antibody staining, selected slices were washed for 3×10min in PBS on rotary shaker at room temperature. The slices were blocked and permeabilized for 30 min (PBS 0,01M / Triton 0,4% / BSA 5%, Sigma Aldrich, USA, Missouri, St. Louis) on rotary shaker. The first antibody (anti-GFAP, PA5-16291, Thermofisher, USA, Massachusetts, Waltham) was applied overnight at 4 degree Celsius (PBS 0,01M / Triton 0,2%). The slices were washed for 3×10min in PBS on rotary shaker at room temperature. The second antibody (Alexa 488 goat anti rabbit, A11034, Thermofisher, USA, Massachusetts, Waltham) was applied for 2 to 3 hours (PBS 0,01M / Triton 0,2%). Finally, the slices were washed for 3×10min in PBS on rotary shaker at room temperature, mounted and stained with DAPI (Vectashield, H-1200-10, USA, California, Burlingame). The slices were imaged with a Zeiss microscope using a 5x objective. The exposure time for DAPI, Alexa 488 and mCherry was 40, 800 and 70 milliseconds respectively. Shading correction was done in Fiji (Based on ImageJ from NIH, USA, Maryland, Bethesda) using the BaSiC plugin ^28^. Stitching was done in Fiji using the Grid/Collection plugin ^29^.

### Statistical procedures

All statistical tests with few samples were done by assuming a normal distribution by means of a double-sided t-test. Statistics for the inhibition latency were calculated by bootstrapping the peri inhibition time histograms of the spikes of sorted units with 1000 repetitions.

---

**Supplementary Figure 1.**
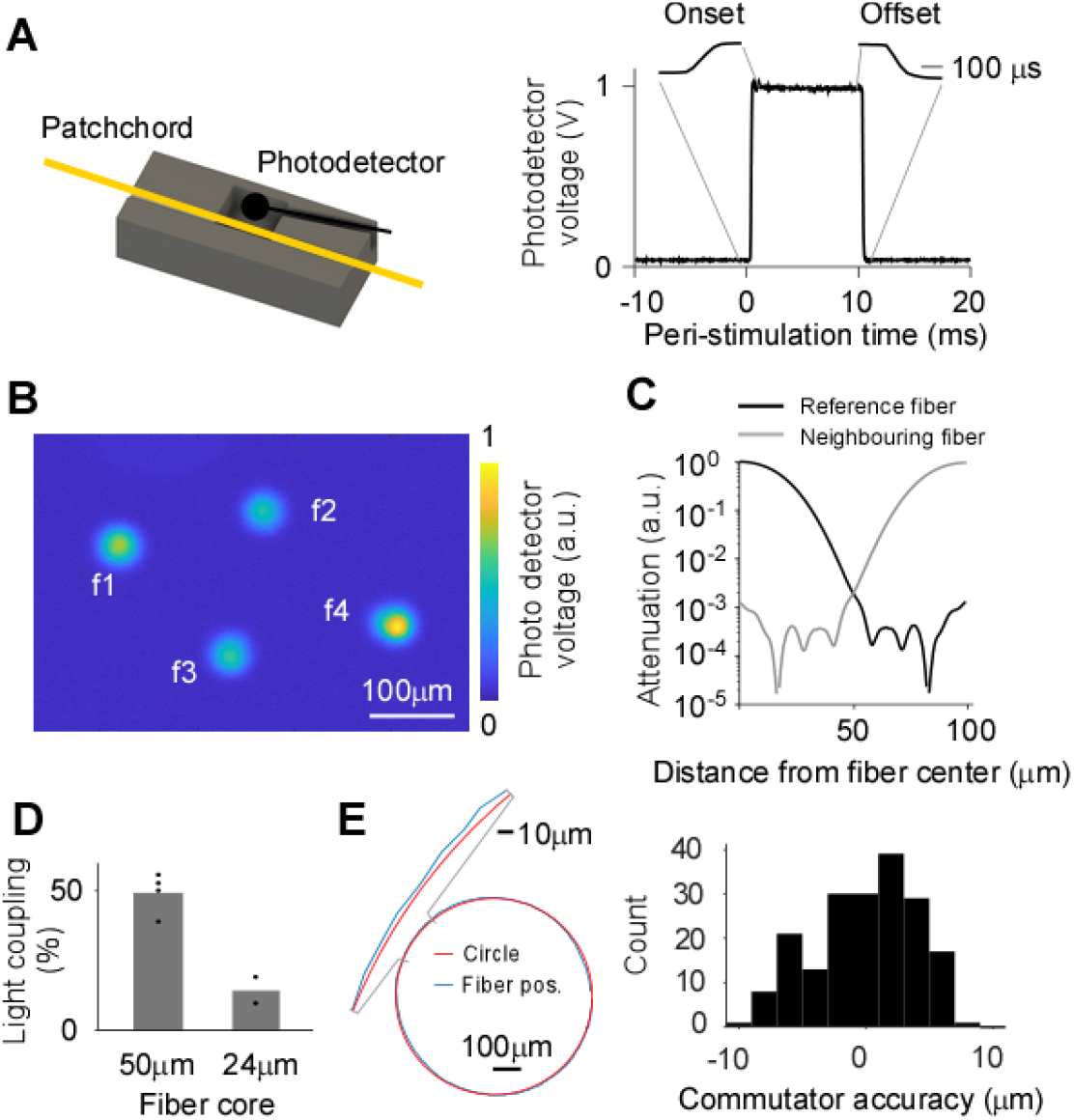
Active patch cord and optical commutator. A: To be able to tune the laser system in the freely moving animal and to automatically find the fiber locations, we attached a linear photodetector to the patch cord. A fraction of the light from the fibers was transmitted through the patch cord furcation tube and detected by the photodiode (left panel). The rapid targeting of the fibers was detected with the photodiode (right panel). B: Example readout from the active patch cord while the galvo scanner mapped four fibers in the ferrule. Scale bar, 100 μm. C: The optical isolation of one fiber (black line) and the mirrored version (gray line) suggests that two fibers with a separation of 100 μm can be individually addressed. D: Input laser power at ferrule divided by output power after passing the patch cord. E: Mechanical stability of the optical commutator. Red circle is the ideal rotation and the blue circle is the center of the optical fiber. Zoomed in section indicates minimal deviations (left). Deviation between ideal position and true fiber position (right).

**Supplementary Figure 2.**
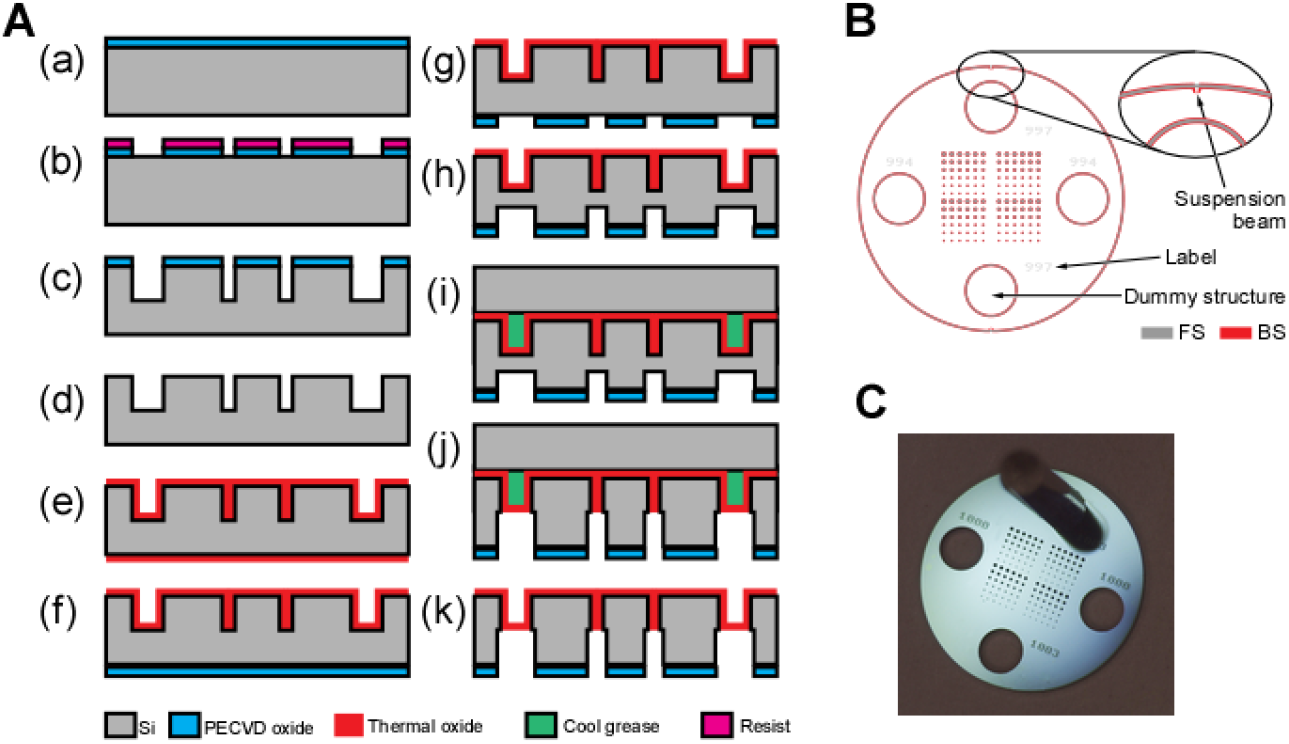
Fiber matrix connector. A: Main processing steps to realize the Si-based connector plates of the fiber connector: (a) Deposition of PECVD SiO_x_ masking layer on wafer front side (FS), (b) patterning of SiO_x_ using lithography and RIE, (c) FS DRIE to an etch depth of 200 μm, (d) wet chemical etching of SiO_x_ layer, (e) thermal oxidation, (f) deposition of 2 μm SiO_x_ in wafer rear side (RS), (g) patterning of RS SiO_x_, (h) DRIE to 150 μm etch depth, (i) wafer fixation on handle wafer using Cool Grease, (j) DRIE etch through, and (k) release from handle wafer. B: The layout of the lithography mask of a connector plate. The gray polygons indicate those areas patterned by deep reactive ion etching from the wafer FS. The etch pattern from the wafer BS was achieved by adding the gray and red polygons. The larger outer ring defined the plate geometry. The structures on the top and bottom indicate beams that suspend the connector plate inside the silicon wafer during the entire fabrication process. Each plate with a diameter of 5 mm comprises two pairs of circular through-holes with equal diameters of ca. 1 mm. One pair of holes will be used for guide-pins and the other pair will be used for anchoring pins. The anchoring pins anchor the plate in the connector sleeve and the fiber bundle.

**Supplementary Figure 3.**
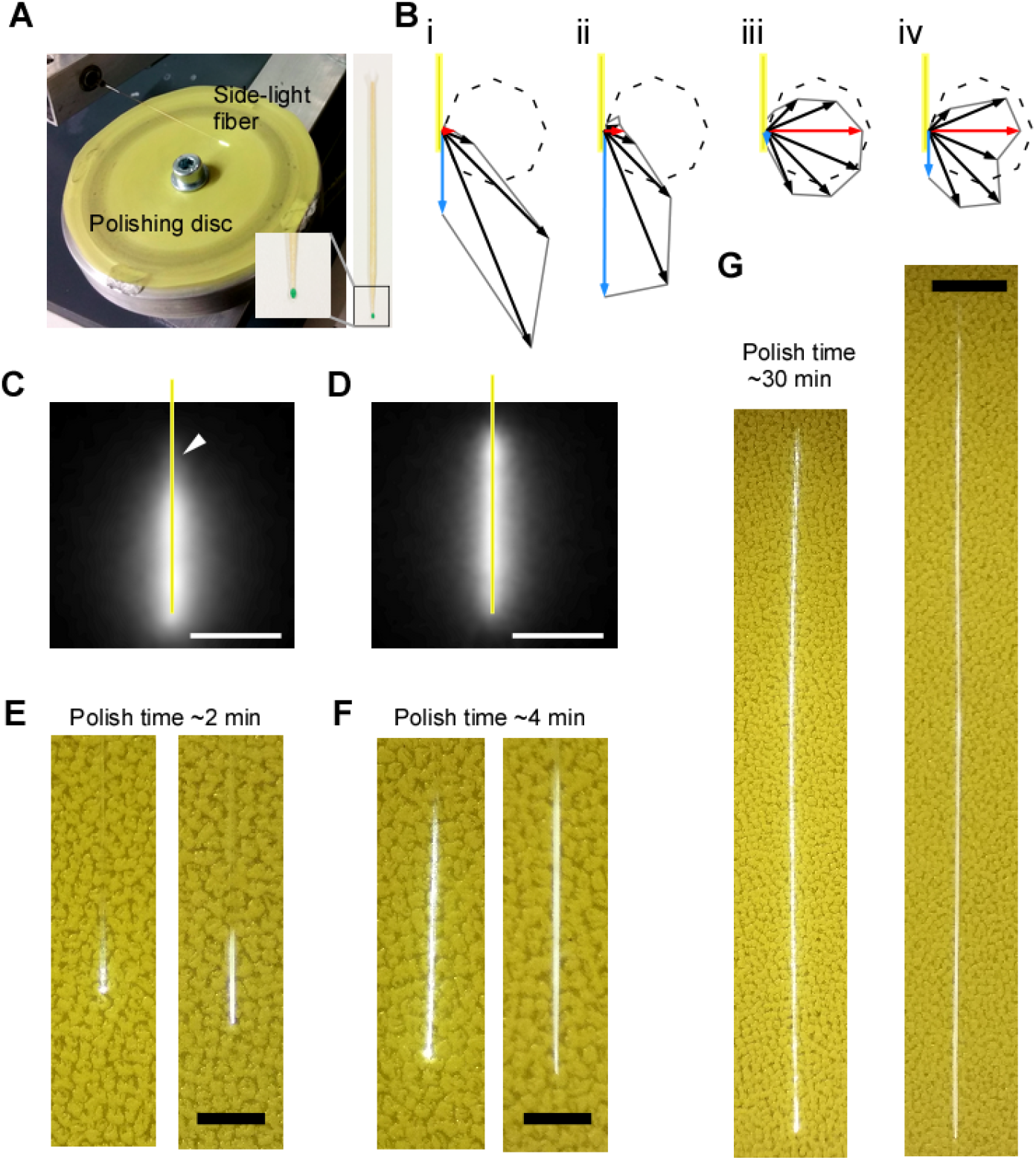
Manufacturing and types of side-emitting optical fibers. A: Polishing plate for manufacturing side-emitting fibers. B: Emission angle of uncoated fiber (i), uncoated fiber glued to a transparent thin film (ii), coated fiber (iii), and coated fiber glued to a transparent thin film (iv) plotted in relation to a Lambertian distribution (dashed circle). Note that the coating enhances the side-emitting component (red arrow) in relation to the forward component (blue arrow). For sideemitting fibers glued onto a transparent thin film laminar electrode, the fiber coating ensures that the light was transmitted through the thin film (red arrow in the fourth panel) rather than to the front (blue arrow in the second panel). Dashed circle describes an ideal Lambertian emitter. C: An uncoated two millimeter fiber in milk. Scale bar, 1 mm. D: A coated two millimeter fiber in milk. Scale bar, 1 mm. E-G: Uncoated (left) and coated (right) fibers polished for a length of 0.5, 2, and 10 mm, respectively. Scale bar, 500 μm, 500 μm, and 1 mm, respectively.

**Supplementary Figure 4.**
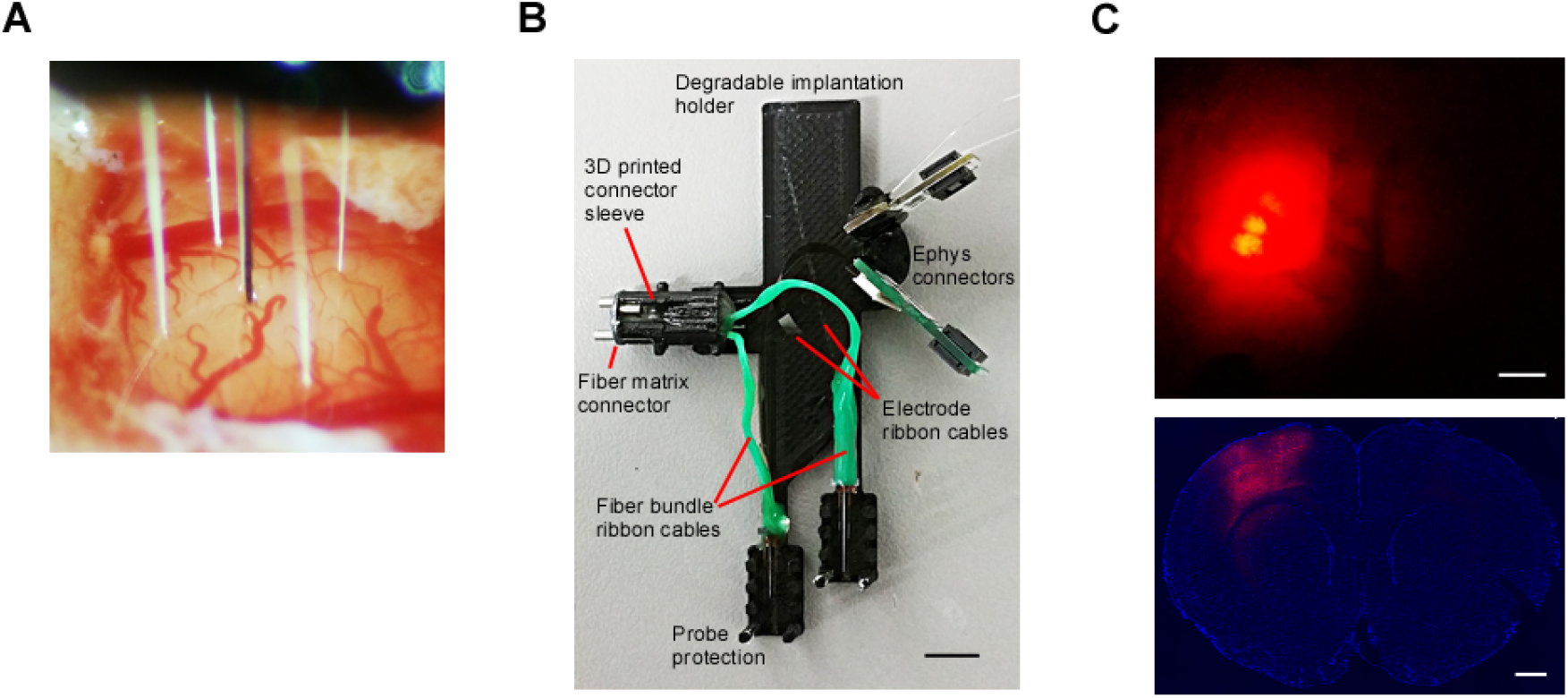
Implantation of multiple fibers and electrodes. A: The brain dimpling was barely detectable when inserting the fibers and electrodes. The BE-probe, consisting of a silicon electrode next to two side-emitting fibers, is surrounded by four additional side-emitting fibers (fiber matrix). B: Temporary implant holder for individual targeting of multiple brain areas with multiple BE-probes and additional fibers. Scale bar, 5 mm. C: In vivo mCherry expression (AAV-hSyn-eNpHR3.0-mCherry-WPRE, 561nm excitation light at 610nm low pass collection filter) was used to guide the implantation (upper panel, view from the top onto the brain’s surface). Corresponding coronal section at bregma 1.2 mm (lower panel). Scale bar, 1 mm.

**Supplementary Figure 5.**
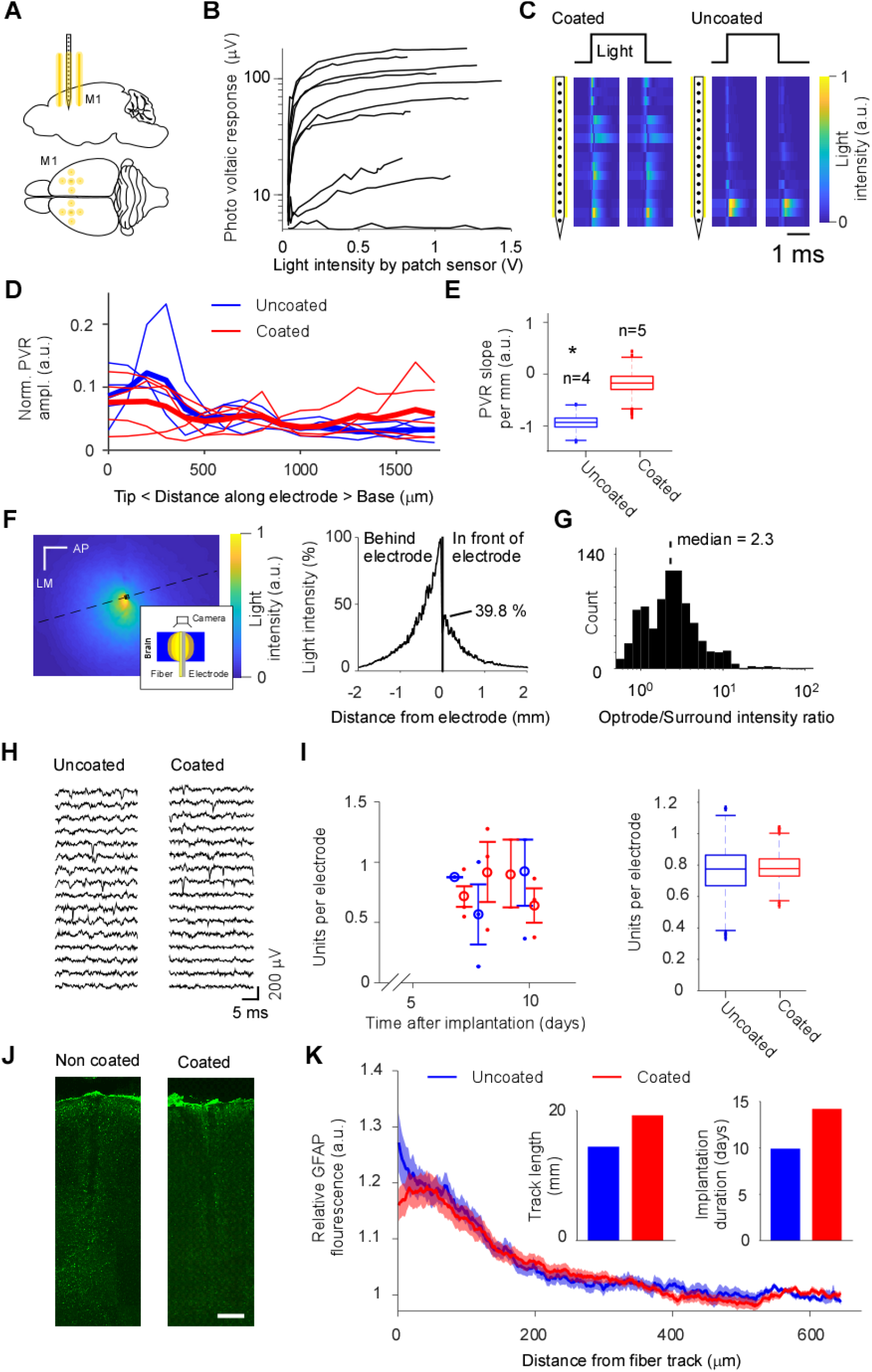
BE-probe performance in the freely moving animal. A: Implanted BE-probes and fiber matrices. B: Assessing the BE-probe functionality using the Photo Voltaic Response (PVR) to make sure that the fibers are intact. C: Example PVR distribution along the electrode for coated and uncoated fiber. D: The PVR along the electrode shank for all coated and uncoated fibers. E: Quantifying the light emission along the electrode for the uncoated and coated fiber by means of the slope of the PVR along the electrode (see panel E). A negative slope is associated with a stronger PVR towards the tip. F: The light distribution of yellow light (589 nm) across the cortical surface from a vertical BE-probe in the motor cortex of a freshly cut mouse brain (left panel). A roughly 5 mm thick transverse slice was put horizontally and was penetrated from below by a BE-probe such that the light distribution could be imaged on the cortical surface from above (left panel, inset). Intensity cross-section (dashed line in the left panel) to compare the light intensity at the electrodes and the fibers (right panel). Scale bar, 0.5 mm. G: Comparing the PVR for BE-probe fibers and fiber matrix. The relative light intensity caused by BE-probe fibers are larger than those of the fiber matrix. H: Examples of extracellular signals for an uncoated fiber and for a coated fiber. I: Quantification of electrophysiology quality as a function of days after implantation and coated (red) versus non-coated (blue) fiber-electrode combinations. J: Representative histological examples for GFAP immunostaining (green) for uncoated and coated fibers after 11 days of implantation. Scale bar, 200 μm. K: Histological quantification of the GFAP signal as a distance to the BE-probe.

**Table 1.** Implanted fibers and electrodes

|  | Optical fibers |  |  |  |  |  |
| --- | --- | --- | --- | --- | --- | --- |
|  | Right hemisphere |  | Left hemisphere |  | Silicon probes |  |
| Animal | Fiber matrix (x4) | BE-probe (x2) | Fiber matrix (x4) | BE-probe (x2) | R.H. | L.H. |
| 540 | NC | NC | C | C | ATLAS | BLBT |
| 541 | C | NC | C | C | ATLAS | BLBT * |
| 542 | C | NC | C | C | ATLAS | BLBT * |
| 548 | C | NC | C | C | ATLAS | BLBT |
| 549 | C | (C) | C | C | ATLAS | BLBT |
| 550 | NC | C | NC | NC | ATLAS | BLBT * |
| 551 | NC | C | NC | NC | ATLAS | BLBT * |

NC=Not coated, C=Coated, R.H. = Right hemisphere, L.H. = Left hemisphere. \* = Large fluctuations (>1mV) in the extracellular signal fluctuations. These fluctuations were most likely due to the reuse of the zif-clip connector and those probes were excluded from further analyses. () = In one BE-probe the fibers were defect since there was no PVR. The ATLAS and BLBT probes are Michigan-style silicon-based intracortical probes. The BLBT probe was realized as a part of BrainLinks-BrainTools.

**Table 2.** Injection, implantation and euthanasian dates

| Animal | Inj. Date | Impl. Date | Sacr. Date | Implantation dur. (days) | Expr. dur. for hist. (days) |
| --- | --- | --- | --- | --- | --- |
| 540 | 20 Dec -19 | 22 Feb -20 | 15 Mar -20 | 22 | 64 |
| 541 | 20 Dec -19 | 27 Feb -20 | 15 Mar -20 | 17 | 70 |
| 542 | 20 Dec -19 | 27 Feb -20 | 15 Mar -20 | 17 | 70 |
| 548 | 23 Jan -20 | 5 Mar -20 | 15 Mar -20 | 10 | 42 |
| 549 | 25 Jan -20 | 5 Mar -20 | 15 Mar -20 | 10 | 40 |
| 550 | 25 Jan -20 | 6 Mar -20 | 15 Mar -20 | 9 | 41 |
| 551 | 27 Jan -20 | 6 Mar -20 | 15 Mar -20 | 9 | 39 |

## Acknowledgements

We would like to thank Ofer Yizhar for comments on an earlier version of this manuscript. This work was supported by the Bernstein Award 2012 (01GQ2301), the cluster of excellence BrainLinks-Brain-Tools (EXC 1086), the Deutsche Forschungsgemeinschaft (DFG) via the grants DI 1908/3-1 and 1908/6-1, as well as the ERC Starting Grant OptoMotorPath (338041), all to I.D, and the Research Innovation Fund to D.E.

## Author contributions

D.E., P.R. and I.D. conceived the study and wrote the manuscript. D.E. designed optical hardware with help from A.T. and A.S.. D.E. performed in-vivo experiments. D.E. and P.R. designed the silicone connector plate. P.R. manufactured the silicone connector plate. D.E. performed data analysis. A.S., M.A. and D.E. perfused animals. D.E. performed the histology with help from M.A. K.S. designed, fabricated, assembled and characterized the BLBT probes. D.E. performed the light measurement in the mouse with help from B.C.

## Data Availability

The data that support the findings of this study are available from the corresponding authors upon reasonable request.

